# Dissemination of influenza B virus to the lower respiratory tract of mice is restricted by the interferon response

**DOI:** 10.1101/2023.10.19.563157

**Authors:** Lara S.U. Schwab, Thi H. T. Do, Marios Koutsakos

**Affiliations:** Department of Microbiology and Immunology, University of Melbourne at The Peter Doherty Institute for Infection and Immunity, Parkville, Victoria, Australia

**Keywords:** influenza, innate immunity, interferon, Mx, upper respiratory tract

## Abstract

The global burden of disease caused by Influenza B virus (IBV) is substantial, however IBVs remain overlooked. Understanding host-pathogen interactions as well as establishing physiologically relevant models of infection are important for the development and assessment of therapeutics and vaccines against IBV. Here, we assessed an upper respiratory tract (URT)-restricted model of mouse IBV infection, comparing it to the conventional administration of virus to the total respiratory tract (TRT). We found that URT infections with different strains of IBV resulted in limited dissemination of IBV to the lungs. Infection of the URT did not result in weight loss or systemic inflammation even at high inoculum doses and despite robust viral replication in the nose. Dissemination of IBV to the lung was enhanced in mice lacking functional type I IFN receptor (IFNAR2) but not IFNγ. Conversely, in mice expressing the IFN-inducible gene Mx1 we found reduced IBV replication in the lung and reduced dissemination of IBV from the URT to the lung. Both URT and TRT inoculation with IBV resulted in seroconversion against IBV. However, priming at the TRT conferred superior protection from a heterologous lethal IBV challenge compared to URT priming, as determined by improved survival rates and reduced viral replication throughout the respiratory tract. Overall, our study establishes a URT-restricted IBV infection model, highlights the critical role of IFNs in limiting dissemination of IBV to the lungs but also demonstrates that the lack of viral replication in the lung may impact protection from subsequent infections.

**Importance:** Our study investigated how IBV spreads from the nose to the lung of mice, the impact this has on disease and protection from re-infection. We found that when applied to the nose only, IBV does not spread very efficiently to the lungs in a process controlled by the interferon response. Priming immunity at the nose only was less protective from re-infection than priming immunity at both the nose and lung. These insights can guide the development of potential therapies targeting the interferon response as well as of intranasal vaccines against IBV.

## Introduction

Influenza A and B viruses (IAV and IBV) cause significant morbidity and mortality every year during annual epidemics. IBV on average accounts for ∼25% of influenza cases, but this can be significantly higher in certain seasons and/or geographic locations(1). IBV disease is typically more severe in school-aged children and adolescents, and can be associated with a variety of systemic complications (2-4). In addition to preventative vaccination, therapeutic interventions exist, namely neuraminidase inhibitors and polymerase inhibitors, to treat influenza infections including IBV. However, these typically need to be administered early after infection for maximum efficacy and could lose effectiveness when resistant virus variants emerge. Additional therapeutics interventions are thus needed to mitigate the clinical disease caused by influenza viruses, especially those that could be administered prophylactically to high-risk groups. The development of such therapeutics requires a thorough understanding of virus-host interactions and the processes that cause severe disease. However, our understanding of these processes in the context of IBV are limited.

The use of animal models can be instrumental, not only in assessing the effectiveness of novel therapeutics but also in determining mechanisms and pathways that underpin their actions. While ferrets are a gold standard for influenza infection, mice offer considerable advantages including lower cost, easier accessibility and easy of genetic manipulation for dissecting mechanisms. However, most studies utilising mice employ direct deposition of a large volume and a high viral dose to the lung, which results in robust infection including considerable weight loss. Although infection of the lower respiratory tract is associated with more severe disease, such methods of inoculation are not necessarily reflective of human infection or disease, whereby infection can be established by a low amount of virus and can often be confined in or start at the upper respiratory tract (URT) prior to disseminating to the lower respiratory tract (LRT). In addition, it is becoming increasingly evident that there are considerable immunological differences between the URT and LRT(5, 6). As such, it is pertinent to understand the virus-host interactions that affect the dissemination of influenza viruses, including IBV, from the URT to the LRT.

The development of URT-restricted infection models for IAV has yielded novel insights into the viral and host factors that affect dissemination of IAV across the respiratory tract, such as the contribution of type I and type III interferon (IFN) in restricting IAV replication(6), as well as into the pre-clinical assessment of novel host-directed therapeutics(7). As these have been exclusively focused on IAV, and whether they apply to IBV in unknown, we used a mouse model to assess the potential of IBV to disseminate from the URT and TRT and factors that underpin restriction of IBV to the URT. Finally, we assessed the potential impact of URT-restricted IBV infection to subsequent immunity in the LRT and protection from challenge. Our data provide novel insights about IBV infection in mice that may inform the development and assessment of more effective therapeutics against IBV.

## Materials & Methods

### Viruses

Influenza A (A/Puerto Rico/8/34 – PR8, A/Udorn/1972) and B (B/Lee/40, B/Victoria/2/1987, B/Yamagata/6/1988, B/Florida/04/2006) viruses were grown in embryonated chicken eggs at 35°C for 3 days. Influenza B (B/Malaysia/2506/04, B/Phuket/3073/2013) were grown in MDCK cells at 33°C for 3 days. Viral titres of all stocks were determined by plaque assay on MDCK cells (ATCC).

### Mice and infections

Female or male C57BL/6L mice aged 6–12 weeks were obtained from the Animal Resources Centre (Perth, Western Australia) or the Biological Research Facility in the Department of Microbiology and Immunology at the University of Melbourne (Melbourne, Victoria). IFNaR2^-/-^, IFNγ^-/-^, and B6.A2G-MX1 mice were bred and maintained at the Biological Research Facility in the Department of Microbiology and Immunology at the University of Melbourne (Melbourne, Victoria). All animal work was conducted in accordance with guidelines set by the University of Melbourne Animal Ethics Committee (ethics approval number 21799). For total respiratory tract (TRT) infection, mice were infected intranasally with 50μl of influenza A or B virus at the indicated doses diluted in PBS under isoflurane anaesthesia. For upper respiratory tract (URT) infection, virus was deposited in 10ul on the nares of mice at the indicated doses diluted in PBS without isoflurane anaesthesia. Blood was collected via the submandibular or cardiac route (terminal).

### Viral titre determination

Lungs and nasal turbinates of mice were homogenized and clarified supernatants were obtained by centrifugation. Lung and nasal tissue samples were stored at -80°C and viral titres were determined by plaque assay in duplicate as previously described (8).

### Cytokine analysis in serum

Cytokines were measured in serum using the Mouse Inflammation Panel LEGENDplex cytometric bead array kit (Biolegend). Data were analysed in FlowJo and analyte concentrations were determined using a sigmoidal 4PL standard curve in Prism (GraphPad).

### Analysis of HA-specific antibodies by ELISA

Antibody binding to influenza B HA proteins was tested by ELISA. The expression of recombinant B/Lee/40 and B/Florida/04/2006 HA proteins has been described previously(9). For ELISA, 96-well Maxisorp plates (Thermo Fisher) were coated overnight at 4°C with 2 μg/ml recombinant HA proteins. After blocking with 1% FCS in phosphate-buffered saline (PBS), duplicate wells of serially diluted plasma were added and incubated for 2 h at room temperature. Plates were washed in PBS-T (0.05% Tween-20 in PBS) and PBS before incubation with 1:10,000 dilution of HRP-conjugated anti-mouse IgG (Sigma) for 1 h at room temperature. Plates were washed and developed using TMB substrate (Sigma), stopped using sulphuric acid and read at 450 nm. Endpoint titers were calculated as the reciprocal serum dilution giving signal 2× background using a fitted curve (4 parameter log regression).

### Statistical analysis

Comparison of unpaired data from 2 groups was performed by an unpaired t test. Comparison of unpaired data from 2 groups was performed by a one-way ANOVA with Tukey’s correction for multiple comparisons or a two-way ANOVA with Sidak’s correction for multiple comparisons. Survival analysis was performed using a Log-rank (Mantel-Cox) test.

## Results

### Limited dissemination of IBV from the upper to the lower respiratory tract in mice

Previous studies have demonstrated variability in the potential of IAV isolates to disseminate from the upper to the lower respiratory tract of mice(6, 10), but such information is lacking for IBV. To address this question, we compared 6 different IBV strains belonging to the Ancestral (B/Lee/40), B/Yamagata (B/Yamagata/16/88; B/Florida/04/2006; B/Phuket/3703/2013) or B/Victoria (B/Victoria/2/87; B/Malaysia/2506/2004) lineages. The IAV PR8 (not disseminating to lung) and A/Udorn/72 (disseminating to lung) were included as positive controls(10). We used an inoculum volume of 10 μL, which has been previously shown to restrict delivery to the upper respiratory tract(6, 7, 10), to deliver a low dose of IBV (100 PFU) to the nose of mice and subsequently assessed viral load in the nose and lungs on day 5 post inoculation. Across all isolates tested, we detected high virus titres in the nasal tissue, but virus was either completely absent or only present at low titres in the lungs of infected animals (3 of 6 animals for B/Victoria/2/87 or 1 of 6 animals for B/Florida/04/2006, respectively) (Figure 1A). Increasing the inoculation dose of B/Victoria/2/87 (10^3^ to up to 10^6^ PFU/10 μL) did not increase dissemination to the LRT (4/8, 3/8, 5/8 or 3/8, respectively) (Figure 1B), with total respiratory tract (TRT) infection (50 μL under light anaesthesia) acting as a positive control. To dissect the kinetics of B/Victoria/2/87 dissemination to the LRT, we applied 100 PFU to the URT or TRT and assessed viral titres up to 9 days after infection both in nasal tissue and lung. Both URT and TRT infections resulted in robust and comparable viral replication in the nasal tissue between day 3-9 post-infection. In contrast, viral titres could only be detected in the lung in a subset of URT-inoculated mice between day 4-7 post infection, while all TRT-inoculated mice showed high viral load between day 3-7 (Fig 1C). We note that throughout these experiments (Figure 1A-C), URT inoculation variably resulted in LRT dissemination in only a subset of mice (average 54.5%, range 37.5-100%). Overall, when IBV is applied to the URT its dissemination to the LRT is limited.

**Figure 1.**
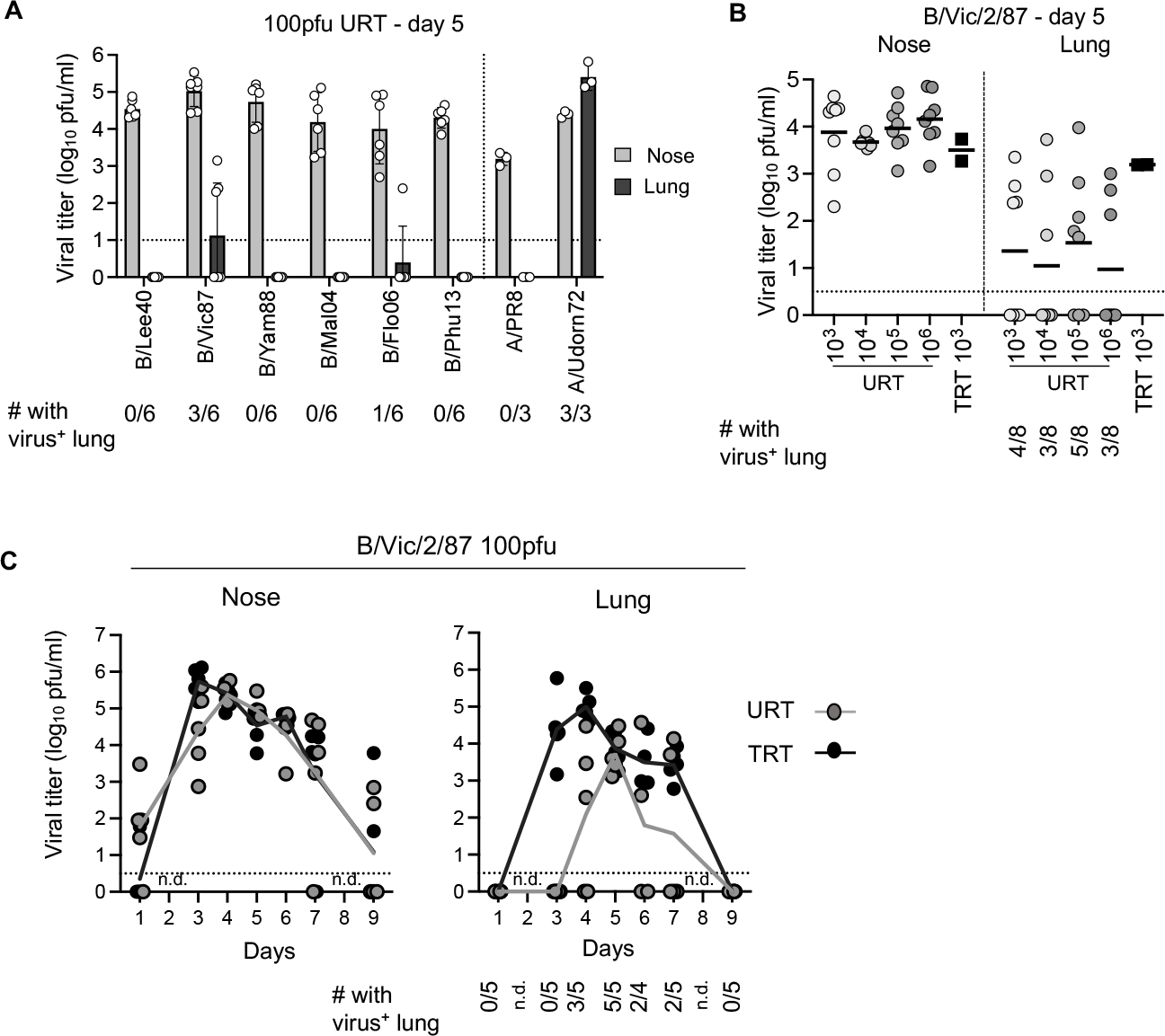
Limited dissemination of IBV from the URT to TRT of mice. **(a)** Mice were infected at the URT with 100PFU of different viruses in 10μl without anesthesia. Viral load was determined in the nose and lung by plaque assay on day 5 post infection. The number of animals in which virus was detected in the lung is shown at the bottom. For IBV n=6 from two independent experiments. A/PR8 and A/Udorn/72 were used as controls (n=3 from one experiment). **(b)** Mice were infected at the URT with B/Vic/87 at different doses in 10ul without anesthesia. Viral load was determined in the nose and lung by plaque assay on day 5 post infection. The number of animals in which virus was detected in the lung is shown at the bottom. Mice infected at the TRT with 10^3^ PFU in 50μl under isoflurane anesthesia were used as positive controls. For URT groups n=8 from two independent experiments, for the TRT group n=2 from one experiment. **(c)** Mice were infected at the URT or TRT (as above) with 100 PFU of B/Vic/87 and viral load was determined in the nose and lung by plaque assay at different timepoints (n=4-5 mice per timepoint).

### IBV upper respiratory tract infection does not cause severe disease in mice

To further characterize URT infection with IBV, we next determined whether URT inoculation could lead to severe disease. We infected mice with a high dose (10^5^ PFU/animal) of B/Lee/40 or B/Florida/04/2006 at either the URT or TRT and assessed weight loss over the first 7 days. As expected TRT infections led to significant weight loss (Fig 2A), with all mice reaching humane endpoint by day 6 post inoculation (Fig 2B). In contrast, URT infection with the same dose did not result in any weight loss or mortality. Consistently, in TRT but not URT-inoculated mice, we could detect inflammatory cytokines like IFNγ, MCP-1 and IL-6 in serum on day 5 post infection, although these varied between the 2 IBV strains studied. This is suggestive of systemic inflammation consistent with the severe disease observed in these mice, but not URT-inoculated mice. These results indicate that although the URT can support high levels of viral replication (Figure 1C), such infections only cause mild disease.

**Figure 2.**
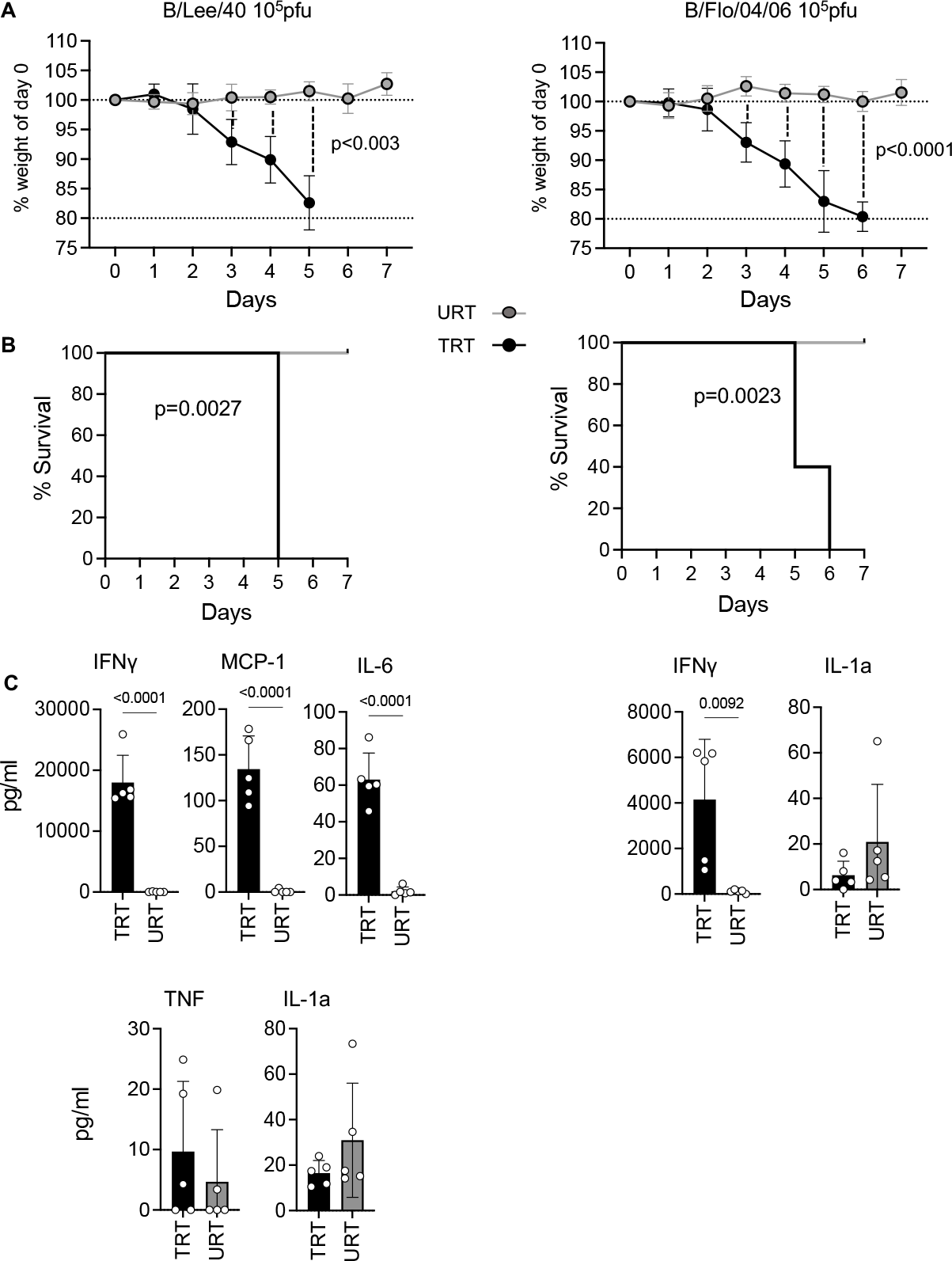
URT IBV infection does not cause severe disease. **(a)** Mice were infected at the URT or TRT with 10^5^ PFU of B/Lee/40 or B/Florida/04/2006 and weight loss was monitored for 7 days. Weight loss at each timepoint was compared using a 2-way ANOVA with with Sidak’s correction for multiple comparisons. **(b)** Survival curves for mice infected at the URT or TRT with 10^5^ PFU of B/Lee/40 or B/Florida/04/2006. The two curves were compared using a Log-rank (Mantel-Cox) test. **(c)** Cytokines were measured in serum on day 5 after URT or TRT infection with 10^5^ PFU of B/Lee/40 or B/Florida/04/2006. The two groups were compared using an unpaired t test. Throughout the figure n=5 mice/group. The two virus strains were tested in independent experiments.

### Type I IFNs but not type II IFNs prevent IBV dissemination from the upper to the respiratory tract

IFNs have an important role in restricting IAV dissemination to the lung(6). To assess their contribution in the context of IBV dissemination, we performed URT inoculations of WT C57BL/6 mice (with functional IFN signalling) and mice which lack type I IFN receptors (IFNAR2 ^-/-^) or type II IFNs (IFNγ ^-/-^) and assessed virus dissemination to the LRT. We chose the B/Florida/04/2006 strain as it exhibited limited LRT dissemination in our initial screen (Figure 1A). Inoculation of the URT with 100 PFU resulted in robust replication in the nasal tissue which was not impacted by the lack of type I IFN signalling nor type II IFNs (Figure 3A). However, we found a significantly greater frequency of IBV dissemination to the LRT of IFNAR2 ^-/-^ mice compared to WT animals (90 % compared to 20 %) (Figure 3B) and significantly enhanced viral load in the LRT (Figure 3A). In contrast, IBV dissemination to the LRT was not impacted by type II IFNs. Overall, these data demonstrate that type I IFNs restrict dissemination of IBV to the LRT.

**Figure 3.**
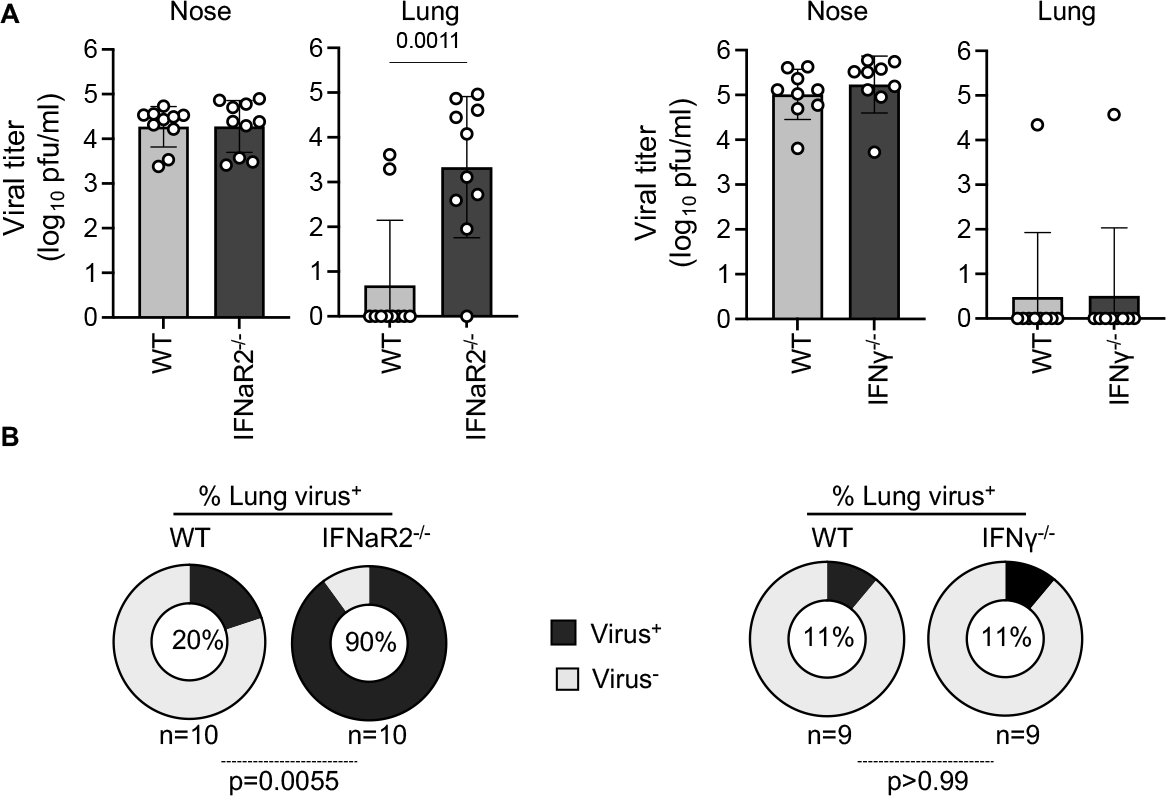
Type I, but not type II IFNs, limit IBV dissemination to the lower respiratory tract. **(a)** WT and sex and age-matched IFNAR1^*-/-*^ or IFNγ^*-/-*^ mice were infected at the URT with 100 PFU of B/Florida/04/2006. Viral load was determined in the nose and lung by plaque assay on day 5 post infection, n=9-10 mice/group from two independent experiments. **(b)** The frequency of mice with detectable virus in the lung on day 5 post-infection is shown for each group. The two groups were compared using a Fisher’s exact test.

### Murine Mx1 restricts IBV dissemination to the lower respiratory tract

To further demonstrate this protective role of IFNs, we assessed whether IFN-stimulated genes (ISGs) could restrict dissemination of IBV to the LRT. Mx1 is a well-described IFN-induced restriction factor for IAV, but little is known about the antiviral activity of Mx1 against IBV. As most laboratory mouse strains, including C57BL/6 mice, lack a functional Mx1 protein(11), we utilised B6.A2G-Mx1 mice, a congenic strain on the C57BL/6 genetic background that expresses a functional Mx1 protein derived from the A2G mouse strain(12). To determine if Mx1 has any antiviral activity against IBV *in vivo*, we firstly performed TRT inoculations of WT C57BL/6 (Mx1-deficient) and B6.A2G-Mx1 mice (with functional Mx1) and determined viral loads in the nasal tissue and the lung on day 5 post inoculation. Following low dose TRT inoculation with 100 PFU of B/Victoria/2/87, we found significantly lower viral titres in the lung of B6.A2G-Mx1 mice (2.385 ± 1.199 log_10_ PFU/ml) compared to WT mice (4.560 ± 0.592 log_10_ PFU/ml) but not in the nasal tissue (Figure 4A).

**Figure 4.**
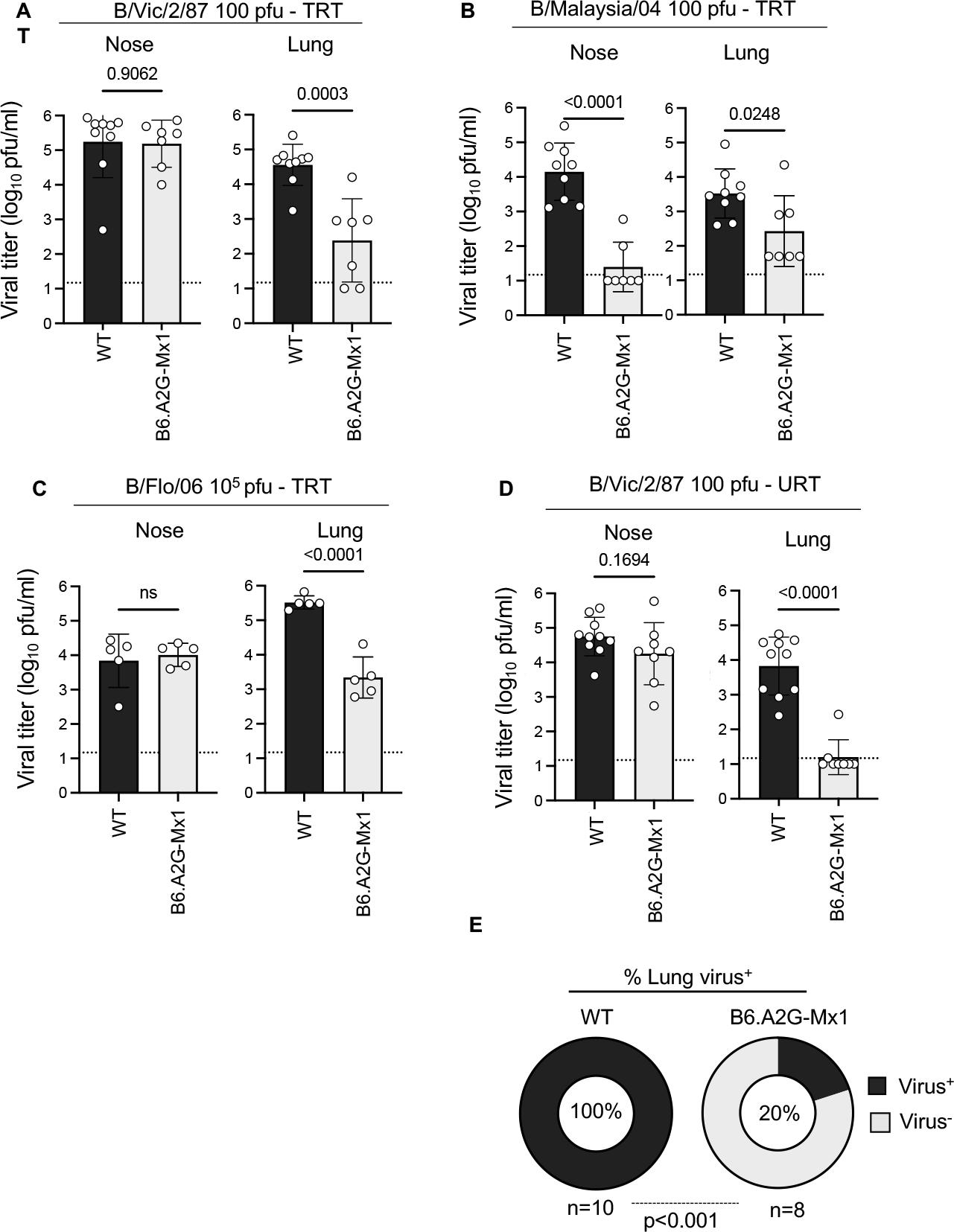
Murine Mx1 limits IBV dissemination to the lower respiratory tract. **(a-c)** C57BL/6 and sex and age-matched A2G-Mx1 mice were infected at the TRT with (a) 100 PFU of B/Victoria/2/87, or 100 PFU of B/Malaysia/2506/2004 (b) or 10^5^ PFU of B/Florida/04/2006 (c). Viral load was determined in the nose and lung by plaque assay on day 5 post infection, n=5-10 mice/group from one (c) or two independent experiments (a, b). **(d)** C57BL/6 and sex and age-matched A2G-Mx1 mice were infected at the URT with 100 PFU of B/Victoria/2/87. Viral load was determined in the nose and lung by plaque assay on day 5 post infection, n=8-10 mice/group, from two independent experiments. **(e)** Frequency of mice with detectable virus in the lung on day 5 post-infection from (d). For (a-d) p-values were determined by an unpaired Student’s t-test. For (e) p-values were determined by a Fisher’s exact test.

As the antiviral effects of Mx1 proteins against IAV are highly virus strain-dependent (13, 14), we tested additional IBV isolates from the B/Victoria lineage (B/Malaysia/2506/2004) or B/Yamagata (B/Florida/04/2006) for their susceptibility to restriction by Mx1. Following low dose TRT inoculation with 100 PFU of B/Malaysia/2506/2004, we found significantly lower viral titres in the lung of B6.A2G-Mx1 mice (2.429 ± 1.027 log_10_ PFU/ml) compared to WT mice (3.523 ± 0.716 log_10_ PFU/ml), as well as in the nasal tissue (B6.A2G-Mx1: 1.397 ± 0.714 log_10_ PFU/ml, WT: 4.149 ± 0.828 log_10_ PFU/ml) (Figure 4B). Mx1-mediated restriction of viral replication in the lung, but not the nose, was also observed after TRT inoculation with a high dose (10^5^ PFU) of B/Florida/04/2006 (B6.A2G-Mx1: 3.340 ± 0.595 log_10_ PFU/ml; WT: 5.515 ± 0.192 log_10_ PFU/ml)(Figure 4C). As we have only assessed viral load on day 5 post infection, it is unclear to what extent an effect of Mx1 on nasal viral load might be evident at other timepoints. Nonetheless, overall, these data demonstrate the potential of Mx1 to restrict IBV replication *in vivo*.

Having established the *in vivo* antiviral effects of Mx1 against IBV, we next tested whether Mx1 could restrict dissemination of IBV to the LRT. We inoculated WT (Mx1-deficient) and B6.A2G-Mx1 mice (with functional Mx1) with 100 PFU of B/Victoria/2/87 at the URT since it was the only isolate capable to disseminating to the LRT in WT mice (Figure 1A). Despite similar viral load in the nasal tissues of WT and B6.A2G-Mx1 mice (Figure 4D), viral dissemination to the lung was only detected in 2/10 B6.A2G-Mx1 mice but in 10/10 of WT mice (Figure 4D and E). These data demonstrate that Mx1 can restrict IBV dissemination to the lower respiratory tract, further supporting a role for IFNs in this protective phenotype.

### Priming of the TRT provides superior protection from severe heterologous challenge

Although the restricted dissemination of IBV to the LRT may limit disease severity, lack of viral replication in the LRT could impact the generation of protective immunity at that site. To test this hypothesis, we primed C57BL/6 mice with 10^2^ PFU of B/Lee/40 at the URT or TRT. At 8 weeks after infection, we collected serum and then challenged the mice with a lethal dose of B/Florida/04/2006 (10^6^ PFU administered to the TRT) and assessed weight loss, survival and virus replication at the upper and lower respiratory tract at 3 days post infection.

At 8 weeks after priming and prior to challenge, IgG antibody titres to homologous and heterologous antigens were measured by ELISA (Fig 5B). Animals primed at the TRT showed significantly higher titres of B/Lee/40 HA antibodies than animals primed at the URT. Interestingly, B/Lee/40 infection resulted in generation of cross-reactive serum antibodies against B/Florida/04/2006 which was significantly higher after TRT infection than after URT infection. Next, animals were challenged with heterologous B/Florida/04/2004. Animals primed with B/Lee/40 showed significantly less weight loss when compared to PBS-treated controls (Fig 5C) and this was reflected in significantly improved survival rates (Fig 5D). Interestingly, priming with B/Lee/40 at the TRT completely protected animals from succumbing to infection whereas URT-primed animals showed 40 % mortality. We then assessed infectious viral titres in upper and lower airways at day 3 post B/Florida/04/2006 infection. We detected virus in the upper respiratory tract of all animals, indicating that no sterilizing immunity was achieved, however both TRT and URT B/Lee/40 priming significantly reduced viral burden in the nose compared to PBS-treated animals (Fig 5E). Importantly, animals primed at the TRT showed reduced levels of virus than URT-primed animals, with 2/5 animals showing complete absence of virus in the lung (Fig 5E). B/Lee/40 priming at the URT did not significantly improve viral burden in the lung compared to PBS-treated animals (Fig 5E). Overall, TRT priming provided superior protection from severe heterologous challenge of the lower respiratory tract.

**Figure 5.**
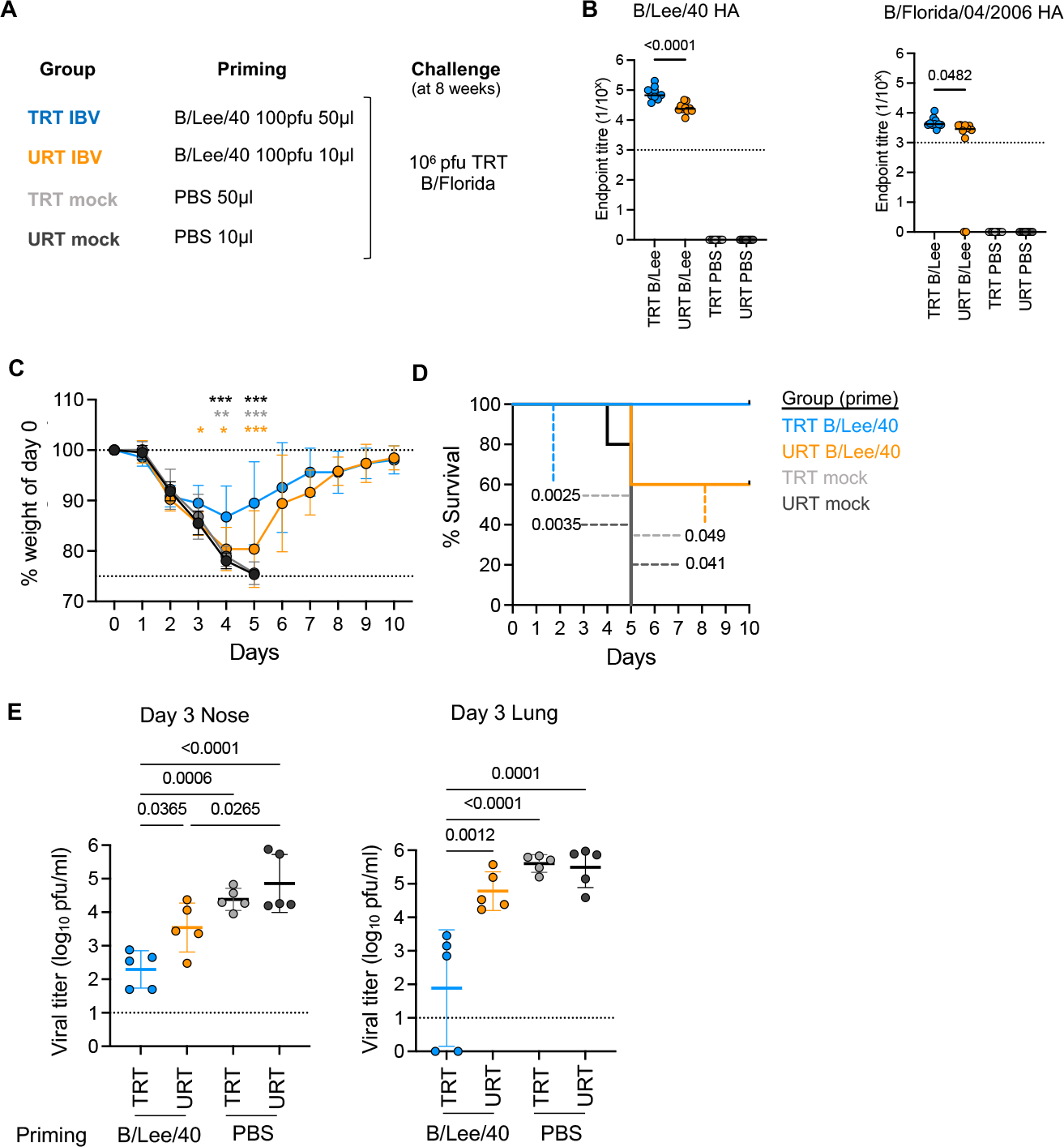
TRT and URT priming provide differential protection from severe heterologous TRT challenge. **(a)** Experimental design. **(b)** At 8 weeks after priming and prior to challenge, mice were bled and serum IgG antibody titres to homologous and heterologous antigens were measured by ELISA. The different groups were compared using a one-way ANOVA with Tukey’s correction for multiple comparisons (n=10 mice/group). **(c)** Weight loss at each timepoint after heterologous TRT challenge. The different groups were compared using a 2-way ANOVA with with Sidak’s correction for multiple comparisons. Statistical significance is shown for URT and PBS groups compared to the TRT group (n=10 mice/group). **(d)** Survival curves after heterologous TRT challenge with B/Florida/04/2006. The two curves were compared using a Log-rank (Mantel-Cox) test. Pairwise-comparisons are color-coded. **(e)** Viral load was determined in the nose and lung by plaque assay on day 3 after heterologous TRT challenge with B/Florida/04/2006. The different groups were compared using a one-way ANOVA with Tukey’s correction for multiple comparisons (n=5 mice/group).

## Discussion

Infections of mice with influenza viruses are usually performed by instilling a large volume with a high infectious dose of IAV or IBV across the TRT of the mouse. While this may ensure robustness of the infection process as well as clinical disease evident by weight loss, these conditions could be considered supraphysiological and not representative of natural infection as the lung environment is overloaded with virus antigen. Alternative models in which virus administration is restricted at the URT have been developed and have provided novel insights into *in vivo* host-pathogen interactions(6, 7, 10, 15, 16). However, these have been limited to IAV and the factors that influence dissemination of IBV to the lower respiratory tract have not been determined. Here, we show that natural IBV dissemination to the lung is limited by the type I IFN response.

Our study shows that IBV does not efficiently disseminate to the lower respiratory tract of mice, with only B/Victoria/2/87 (1/6 isolates tested from the three lineages) showing detectable virus in the lung after URT administration. This observation is consistent with limited dissemination of IBV seen in the ferret model(17). Even for B/Victoria/2/87, dissemination to the lungs was highly variable between experiments and could not be improved by increasing the amount of virus administered to the URT. It will be interesting to determine the virological factors that determine the increased propensity of B/Victoria/2/87 to disseminate to the lower respiratory tract compared to other isolates. We note that the strains tested in our study are human isolates and not mouse-adapted IBV strains which may behave differently. For IAV, dissemination to the lung has been associated with the ability of the NA to overcome inhibitors present in mouse saliva (15). While differences in the IBV NA may similarly impact dissemination to the lung, differences in BNS1 may also contribute given the prominent role of type I IFNs we observed in restricting IBV dissemination. Furthermore, in our experiments, administration of 10^5^ PFU to the URT, which is lethal when administered to the TRT, did induce weight loss or systemic inflammation. Thus, IBV infection of URT of mice may be a useful animal model to understand subclinical disease and the factors that regulate disease tolerance in URT.

The robust dissemination of IBV to the lungs of mice lacking IFNAR2 suggests a strong contribution of type I IFNs in restricting IBV to the URT. Interestingly, we did not observe increased viral replication in IFNAR2^-/-^ mice suggesting type I IFNs may have a lesser role in the URT, although additional experiments are needed to elucidate this question. Indeed, in the context of IAV, replication in the URT is most strongly impacted by type III than type I IFNs (6, 18). Similarly, type III IFNs can also restrict IAV dissemination to the lung(6), which we could not assess in our study as type III IFN deficient mice were not available. Further dissecting the relative contributions of type I and III IFNs can inform the development of host-directed antiviral strategies against respiratory viruses including IBV. Indeed, our data strongly support the notion that antiviral therapeutics that leverage the IFN-response could contribute to protection by limiting dissemination of virus to the LRT.

While previous studies have established the effects of human MxA(19) or mouse Mx1(20) proteins against human and avian IAVs, our study demonstrates an *in vivo* role for murine Mx1 in restricting IBV replication. This is consistent with previous reports of reduced IBV polymerase activity in the presence of Mx1 in an *in vitro* minireplicon system(21). Our experiments further show that Mx-mediated suppression may contribute to restriction of IBV to the URT. It remains pertinent to confirm the role of human MxA protein against IBV as well as to the understand the molecular details of how Mx proteins act against IBV, whether these differ from IAV, and the basis of potentially differential susceptibility of IBV strains to Mx-restriction. Further understanding the role of Mx proteins in restricting IBV replication could inform determinants of susceptibility as well as the development of antiviral therapeutics that leverage the host response.

Although restriction of viral replication in the URT may result in subclinical disease, the lack of viral replication in the lung may impact the generation of local adaptive immunity and subsequent protection. Indeed, we found that URT priming was less protective than TRT priming against a heterologous lethal challenge. Further experiments are needed to determine if differential protection (in the form of reduced viral replication) will be afforded in the context of lower infectious dose or dissemination from the URT to the LRT. Indeed, we found that as few as 100 PFU of IBV are sufficient to establish robust viral replication (Fig 1C, Fig4), even in the absence of weight loss, and it will be interesting to assess protection by URT and TRT priming against such a low dose challenge. It will also be interesting to assess how adaptive immunity in the URT may impact dissemination of IBV to the lung as has been shown for IAV(16). With regards to protection, we hypothesize that URT infection results in lower levels of immunity overall or does not result in the generation of lung tissue-resident B and T cells, thus conferring less effective protection. Consistently, the generation of such mucosal immunological memory after IAV infection of mice is dependent on local inflammatory signals as well as the presence of antigen(22, 23). Further understanding how URT versus TRT priming impact the generation of immunity systemically and in the lung as well as subsequent protection will be critical for the development of effective mucosal vaccines, whose administration is currently often limited to the URT.

Overall, we demonstrate how IBV dissemination to the lower respiratory tract is limited the type I IFN response in mice, which may have implications for understanding IBV pathogenesis. Furthermore, we demonstrate differential protection afforded by URT or TRT priming. These findings may aid the development and assessment of therapeutic or prophylactic interventions against IBV or other respiratory viruses to limit their clinical burden.

## Acknowledgements

The work has been generously supported by the Morningside Foundation and by the Australian National Health and Medical Research Council Investigator grants.

## Author contributions

M.K. designed and supervised the study. L.S.U.S, T.H.T.D. and MK performed experiments and analysed data. L.S.U.S and M.K. drafted the manuscript. All authors revised the manuscript.

